# A preliminary study of HMGB1 defense in Crohn’s disease

**DOI:** 10.64898/2026.07.09.737505

**Authors:** Anne-Marie C. Overstreet, Kylee A. Hunterx, Shivani Patel, Hridai Dharan, Steven Overend, Jeannette S. Messer

## Abstract

**Introduction:** Intestinal barrier failure is a key characteristic of both kinds of inflammatory bowel disease (IBD), ulcerative colitis (UC) and Crohn’s disease (CD). The intestinal barrier, or gut mucosal barrier, is composed of mucus and a tightly interconnected single layer of intestinal epithelial cells (IEC) that line the gut. Together, these components work to contain the gut microbiota within the gut lumen and when they fail, microbes, microbial products, or microbial components cause damage, inflammation, and immune activation in host tissue. We previously reported that High mobility group box 1 (HMGB1) in colonic mucus aggregates bacteria, limits bacterial invasion through mucus, and prevents bacteria from adhering to host tissue. Epithelial surface-associated HMGB1 is decreased in active UC lesions and low levels of HMGB1 are associated with high levels of tissue-adherent bacteria expressing adhesins carrying the molecular target of HMGB1 (ToH1). The study reported here was designed to determine whether HMGB1 defense is also compromised in active lesions from CD patients.

**Methods:** Immunofluorescence microscopy was used to visualize mucus and HMGB1 in tissue from colonic resections performed in CD and non-IBD control patients.

**Results:** Active CD lesions had areas where the IEC were absent or pulling away from underlying tissue along with areas of increased mucus thickness and goblet cells full of mucus highly positive for alpha-linked-fucose residues. The surface associated HMGB1 was also decreased in active CD lesions.

**Conclusion:** Tissue from CD patients exhibited cellular and acellular intestinal barrier defects in comparison to control patients. We observed the previously reported loss of IEC barrier integrity and abnormalities in the amount and distribution of mucus in CD lesions. We also report for the first time that CD is associated with decreased HMGB1 defense at the colon surface.

## Introduction

The inflammatory bowel diseases (IBD), ulcerative colitis (UC) and Crohn’s disease (CD), are diseases of intestinal barrier failure.[1] The intestinal barrier, also referred to as the gut mucosal barrier, is a complex structure that physically separates the microbiota within the intestinal lumen from the host tissues.[2] At the interface with bacteria in the gut lumen, the colonic barrier is composed of mucus that chemically and physically limits contact between microbes and host tissues.[3] Under mucus lies a single layer of tightly interconnected intestinal epithelial cells (IEC).[4] The IEC are all generated from stem cells at the base of colonic crypts and take on specialized functions as they migrate toward the gut lumen.[5] The bulk of the IEC population in the colon is absorptive enterocytes, with smaller numbers of other specialized cells, including mucus-secreting goblet cells. Together, the IEC create a physical barrier against passage of live microbes, microbial products, and microbial components into deeper host tissues.[6] The IEC also produce the components of the acellular barrier and secrete them into the gut lumen.[7] When the intestinal barrier fails, microbes, microbial products, and/or microbial components come into contact with host tissues.[8] This leads to IEC damage, loss of barrier integrity, inflammation, and immune activation. These are the hallmarks of IBD and a growing body of literature has demonstrated that intestinal barrier integrity is the most important indicator of disease progression in IBD patients.[2, 9, 10] However, the pathophysiologic details of how and why the intestinal barrier fails in IBD remain unclear.

Mucus is a complex mixture of water, proteins, small molecules and other components that creates a zone of microbial exclusion between bacteria in the gut and the intestinal epithelium.[11, 12] This relatively sterile layer of mucus is vital to gut health since without it, bacteria adhere to the host epithelium and cause colitis.[12] Mucins are the most abundant of the mucus-associated proteins with Mucin 2 being the most abundant mucin in the colon.[12] Mucin 2 is produced and secreted by goblet cells in the colon, but enterocytes also produce and release components of the colonic mucus, including High Mobility Group Box-1 (HMGB1).[12-14] Both HMGB1 and Mucin 2 play important roles in host defense against bacterial adhesion to host tissues.[15] Mucin 2 is highly glycosylated and polymerizes, contributing to the gel-like nature of mucus.[3] These characteristics create physical resistance to bacterial invasion through mucus, provide foodstuffs that disincentivize virulence, and act as decoy adhesion sites for bacteria.[3] HMGB1 limits bacterial movement through mucus and prevents bacterial attachment to host cells through binding to a small amino acid motif, the target of HMGB1 (ToH1), in bacterial adhesins.[13] When commensal bacteria encounter HMGB1, they down-regulate ToH1+ adhesins, suggesting that HMGB1 also promotes bacterial commensalism in the gut.

We previously reported that HMGB1 is deficient in the colonic mucus of active UC lesions and that HMGB1 levels at the epithelial surface correlate with the numbers of tissue-adherent bacteria expressing ToH1 positive (ToH1+) proteins.[13] As HMGB1 levels decrease, the number of adherent ToH1+ bacteria increases. Mice conditionally deficient in IEC HMGB1 are highly susceptible to colitis (dextran sodium sulfate and IL-10^-/-^ models) and have increased numbers of ToH1+ bacteria adherent to their colonic tissue, even in the absence of challenge.[13, 16] Likewise, bacteria are much more readily adherent to IEC lacking HMGB1 than wild type cells and HMGB1 protects IEC from bacterial damage *in vitro*. While we focused on UC in our initial work, CD is also a disease of intestinal barrier failure and bacterial adhesion to intestinal tissue. This led us to hypothesize that HMGB1 defense is deficient in CD lesions. The goal of this study was to characterize HMGB1 defense in CD lesions to understand how this defense fits into the overall context of acellular mucosal defense in active CD.

## Methods

### Human tissue

The study protocol was approved by the Cleveland Clinic Institutional Review Board. In short, residual, full-thickness, freshly resected tissue was obtained from subjects with Crohn’s disease and controls undergoing colonic resection for clinical management of disease. Control tissues were obtained from disease-free margins of individuals with diseases other than IBD, as previously described.[17, 18] Control tissues were confirmed to have histologically normal epithelium (by hematoxylin and eosin staining) before inclusion in the study. Inflammation was scored by a clinical pathologist based on tissue submitted as part of patient care. Further information about study subjects is available in Table 1.

**Table 1:** Patient characteristics.

| **Demographics** | **Non-IBD** | **CD** |
| --- | --- | --- |
| Total | 12 | 9 |
| Age, mean, (SD), [range] | 57.8 (±14.6) [34-75] | 50.2 (±17.0) [30-76] |
| Age in males | 54.9 (±16.1) [34-75] | 53.8 (±17.6) [30-76] |
| Age in females | 61.8 (±10.8) [45-72] | 45.8 (±15.2) [36-72] |
| **Sex, n (%)** |  |  |
| Male | 7 (58.3%) | 5 (55.6%) |
| Female | 5 (41.7%) | 4 (44.4%) |
| **Disease, n (%)** |  |  |
| Fecal Incontinence | 1 (8.3%) | - |
| Constipation | 1 (8.3%) | - |
| Diverticulitis | 8 (66.7%) | - |
| Cancer | 2 (16.7%) | - |
| **Inflammation Severity** |  |  |
| 1 (none/mild), n (%) | 12 (100%) | 1 (11.1%) |
| 2 (moderate), n (%) | 0 | 3 (33.3%) |
| 3 (severe), n (%) | 0 | 5 (55.6%) |
| **Ethnicity** |  |  |
| White | 10 (83.3%) | 8 (88.9%) |
| Black | 1 (8.3%) | 0 |
| Unknown | 1 (8.3%) | 1 (11.1%) |

### Immunostaining for multiplex immunofluorescence immunohistochemistry (mIFplex-IHC)

Patient tissues were fixed in Methyl-Carnoy’s fixative (60% methanol, 30% chloroform,10% glacial acetic acid) for 3 hours at 4°C and then transferred to 70% ethanol and submitted to the Cleveland Clinic Tissue Histology Core for embedding and sectioning. Paraffin-embedded tissue sections were deparaffinized and equilibrated with water (as follows). Slides were heated for 30 minutes at 60°C, then placed in xylene for 5 minutes (3 times), dunked 15 times in 95% ethanol (2 times), dunked 15 times in 70% ethanol (1 time), placed in water for 1 minute, then placed in TBS-T for 5 minutes (2 times). Slides underwent antigen retrieval with 10 mM sodium citrate + 0.05% Tween 20, pH 6.0 in a steamer for 20 minutes and then were cooled at room temperature for 1 hour. Finally, slides were placed in in TBS-T for 5 minutes (2 times). Each step was performed the indicated number of times, moving the slides into a new container of solution each time the step was repeated.

Tissues were blocked with Superior™ Blocking Buffer at room temperature for 1 hour then stained with 0.17 μg/mL anti-HMGB1 antibody (RRID: AB_1603373) and 10 μg/mL anti-E-cadherin antibody (RRID:AB_355568) diluted in blocking buffer, overnight at 4°C. The next day the slides were placed in TBS-T for 5 minutes (3 times). Secondary antibodies were diluted to a concentration of 2 μg/mL (Alexa Fluor 647 donkey anti-rabbit IgG (RRID:AB_2492288)) or 7.5 μg/mL (Alexa Fluor 555 donkey anti-goat IgG (RRID:AB_3095465)) in blocking buffer and incubated at room temperature for 30 minutes. Slides were placed in TBS-T for 5 minutes (3 times) and then stained with 10 μg/mL Dylight 594 conjugated *Ulex Europaeus* Agglutinin I (UEA I) for 30 minutes at room temperature. Slides were placed in TBS-T for 5 minutes (3 times), counterstained using 10 μg/mL bis-Benzimide H 33258 (Hoechst) dissolved in TBS for 20 minutes in the dark at room temperature, washed with water for 5 minutes, and then mounted with Prolong− glass antifade. Slides were dried at room temperature for 24 hours before imaging.

### Tissue Imaging

Images were captured by microscopists blinded to patient status.

### Whole slide scanning of mIFplex-ICH

Imaging mIFplex-IHC sample slides were scanned using the Vectra PhenoImager HT Automated Quantitative Pathology Imaging System using the PhenoImager HT version 2.1.0 software (Quanterix, Billerica MA). Whole slides were scanned at 20x magnification using the appropriate dichroic filters and spectral unmixing library algorithm generated with Phenochart software v2.2.0 and Inform software v3.1.0 to spectrally unmix the scans. Scale bars are 50 µm.

### Widefield fluorescence microscopy

HMGB1 in colonic mucus was visualized at 40X magnification on the BioTek Lionheart FX Automated Microscope (Agilent). All images were captured using the DAPI (Hoechst) and Cy5 (HMGB1) channels, and images were viewed and edited in the Gen5 3.18 software.

## Results

### Evidence of intestinal barrier abnormalities in CD lesions

We previously demonstrated that HMGB1 released from IEC accumulates in colonic mucus and contributes to acellular defense against bacterial adhesion in the colon.[13] In UC patients, we found little to no HMGB1 at the epithelial surface in active lesions. Alteration of the epithelial surface-associated mucus layer is a key feature of both UC and CD. Previous studies have established that UC is characterized by a thin, often absent layer of mucus.[19] In contrast, CD is characterized by thickened mucus with changes in mucin glycosylation and altered viscoelastic properties.[20, 21] We obtained colonic resections from CD and non-IBD control patients and fixed them in Methyl-Carnoy’s solution to visualize mucus (shown in Table 1).[22] We then characterized mucus using immunofluorescence (IF) staining with the lectin *Ulex Europaeus* Agglutinin I (UEA I). This lectin binds to α-linked-fucose residues, which cap both O- and N-glycans, to label mucus through its glycosylation.[23]

Surgical resection for CD is not curative for the disease, so it is usually performed due to fibrosis and stricture formation, fistulas, or abscesses.[24] Therefore, it was not surprising that the tissue from CD patients was much more friable and exhibited very different architecture than that of control patients. Surface-associated epithelial cells anchored to the underlying lamina propria were present in all of the contol tissues. In contrast, all of the CD tissues had areas that lacked the surface layer of epithelial cells and some sections had no identifiable areas of intact epithelium. Similarly, the epithelium could be seen pulling away from the lamina propria as sheets in many of the CD patients (show in Fig. 1A). As previously reported, many CD patients had areas in which the surface-associated mucus appearted to be thicker than that of controls, but this was highly variable across fields and areas without intact epithelium were devoid of UEA-1 staining. Most strikingly, the majority of the CD patients (all but one in those that had areas of intact epithelium) exhibited intense UEA-1 staining within what appeared to be goblet cells, consistent with abnormality in mucus production, mucus secretion, and/or mucus modification (shown in Fig. 1B). Together, these findings corroborated reports from other studies that the epithelial and mucus barriers of the colon are compromised in CD lesions.

**Fig. 1.**
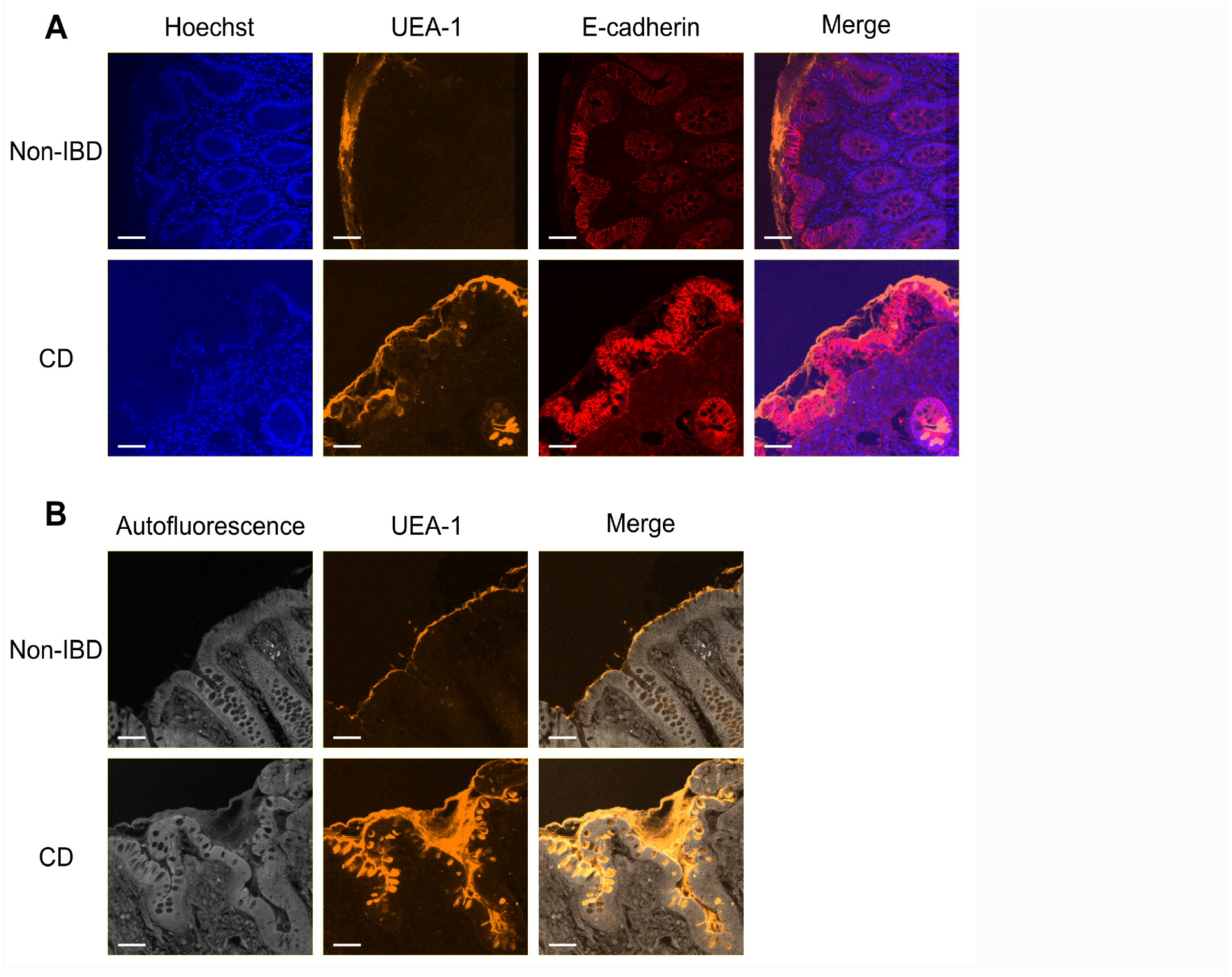
Evidence of intestinal barrier abnormalities in CD lesions. A) IF staining for UEA-1 (orange), E-cadherin (red), and Hoechst (blue). B) IF staining for UEA-1 (orange). Autofluorescence (grey) used to visualize tissue. For both A and B images are representative of n= 9 CD/12 Control patients. Scale bars 50 µm and original magnification 200x.

**Fig. 2.**
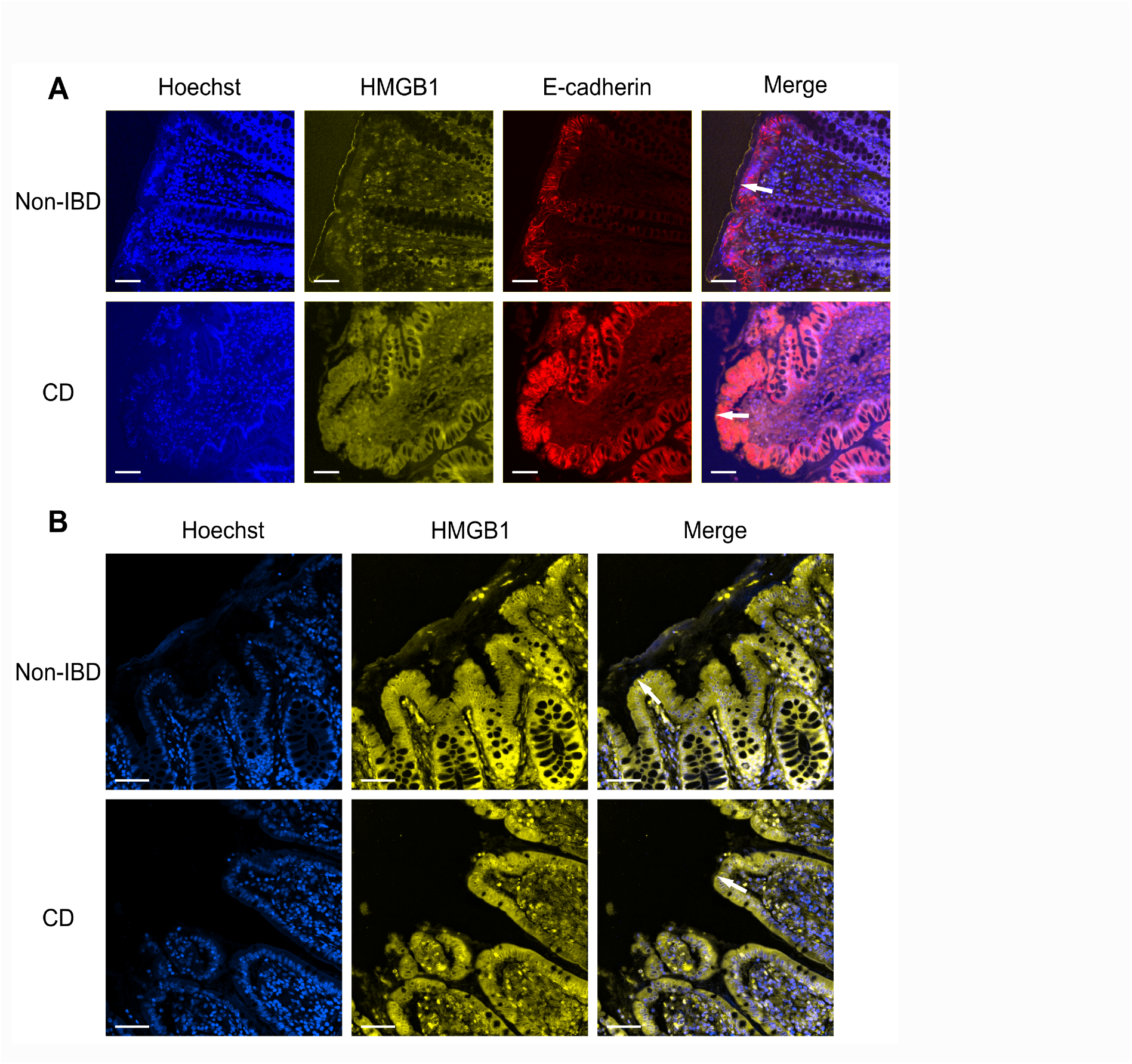
Surface-associated HMGB1 is compromised in CD lesions. A) IF staining for HMGB1 (yellow), E-cadherin (red), and Hoechst (blue). Scale bars 50 µm and original magnification 200x. B) IF stainingfor HMGB1 (yellow) and Hoechst (blue) with images optimized for visualization of the extracellular HMGB1 at the intestinal surface. Scale bars 50 µm and original magnification 400x. For both A and B arrows note the surface of the intestinal epithelum and images are representative of n= 9 CD/12 Control patients.

### Surface-associated HMGB1 is compromised in CD lesions

We next evaluated HMGB1 in the mucus from CD and control tissues. As we previously reported, a layer of HMGB1 associated with the epithelial surface was visible in all of the control tissues.[13] In contrast, HMGB1 was absent in the CD tissues wherever the tissue lacked an intact epithelium. In areas with distortion of the tissue architecture, where epithelial cells were still present, it was possible to find HMGB1 on the surface of the tissue (shown in Fig. 1A, B). However, this was most often a patchy, discontinuous layer of HMGB1. Most interestingly, we did not see accumulation of HMGB1 within areas of what appeared to be thickened mucus, based on UEA-1 staining. Thus, HMGB1 defense is compromised in CD lesions, similar to what was previously reported in UC lesions.

## Discussion

Our findings confirm previously reported mucus abnormalities within CD lesions and provide the first evidence that HMGB1 mucosal defense is also compromised in CD. HMGB1 concentrated in colonic mucus defends the underlying intestinal epithelium against bacterial adhesion and damage that can lead to epithelial barrier failure and immune activation.[13] In the mucus, HMGB1 aggregates bacteria and impairs their movement to prevent bacteria from ever reaching the epithelium. HMGB1 also directly prevents bacterial adhesion to IEC through binding to ToH1 in surface-expressed adhesins. HMGB1 binding to ToH1 blocks adhesin capture of carbohydrate ligands on the surface of host cells, so bacteria do not adhere to the intestinal epithelium. Exposure to HMGB1 also causes commensal bacteria to down-regulate ToH1 containing adhesins, implying that it promotes commensal states in bacteria.

Our discovery that HMGB1 is not uniformly concentrated along the intestinal epithelium in CD lesions suggests that the failure of this component of the acellular colonic barrier contributes to disease in CD. Bacterial adhesion is a well-established feature of CD.[25] Adherent and invasive *E. coli* (AIEC) are perhaps the best characterized of the adherent bacteria linked to CD.[26, 27] These microorganisms use Type 1 fimbrial adhesion, in which the canonical ToH1 positive (ToH1+) adhesin, FimH, anchors bacteria to IEC.[22, 27] In our previous study, we established that the number of adherent bacteria expressing FimH in UC tissue increased as levels of surface-associated HMGB1 decreased.[13] We suspect that low levels of HMGB1 are also contributing to the increased *E. coli* adhesion reported in CD. However, when we attempted immunostaining for FimH in CD tissues, the required staining procedure led to degradation of many of the CD tissue samples and we did not feel that we could confidently evaluate FimH under these conditions.

A defect in HMGB1 was not the only abnormality in mucosal barrier defense that we identified in CD tissues. As previously reported, the cellular barrier composed of IEC was also compromised in CD lesions. Also consistent with previous studies we found that the surface-associated mucus was thicker in many areas of tissue obtained from CD patients. The surface-associated mucus in CD tissue was highly stained by UEA-1 and we saw highly UEA-1 positive cells in the epithelium that were likely goblet cells. Goblet cell secretion defects have been described in CD, so it is possible that these cells are engorged with mucus that is accumulating within the goblet cells.[22] However, we cannot rule out increased mucus production or an alteration in mucus modifications that contribute to the differences in CD mucus. In support of this, other studies have identified differences in lectin staining and glycosylation of mucus in patients with IBD.[28-31] It was very interesting that while mucus thickness was increased in CD, the amount of HMGB1 in the mucus was decreased. This discordance between the two components of mucus may be due to the fact that they are produced in different cell types, the mucins are produced in goblet cells whereas HMGB1 is produced in enterocytes.

Together, our data demonstrate that like UC lesions, CD lesions exhibit defects in HMGB1 defense against bacterial adhesion. However, we do not yet understand whether the defect in HMGB1 defense is a cause or effect of disease or how it relates to other changes in mucus seen in CD. Limitations related to the small number of patients available and the type of tissue needed for this study also restricted our ability to establish a clear relationship between adherent bacteria and defects in HMGB1 defense in CD. Although, previous work in animal and cellular models as well as UC tissues are highly suggestive that the same relationship of increased bacterial adhesion in the absence of surface-associated HMGB1 exists in CD. If true, therapeutics based on restoring or replacing failed HMGB1 defense could have applications in both types of inflammatory bowel disease. This therapeutic strategy has the potential to promote epithelial barrier restoration without harming commensal bacteria, two major goals of IBD therapy.

## Statements

## Acknowledgement

This work utilized resources and equipment provided through the Cleveland DDRCC (NIDDK P30 DK097948) and the Cleveland Clinic Light Microscopy Core. We also thank John Peterson from the Cleveland Clinic Light Microscopy Core for assistance with image capture.

## Statement of Ethics

This study protocol was reviewed and approved by the Cleveland Clinic Institutional Review Board. De-identified tissues were obtained from the Cleveland DDRCC Biobank under an exemption from requiring informed consent.

## Author Contributions

Conceptualization, J.S.M.; methodology, J.S.M., A.-M.C.O., K.A.H; investigation, A.-M.C.O., K.A.H, H.D., S.O., and S.P.; funding acquisition, J.S.M.; project administration, J.S.M.; supervision, J.S.M., A.-M.C.O., and K.A.H.; writing – original draft, J.S.M., A.-M.C.O., and H.D..; writing – review & editing, all authors.

## Data Availability Statement

All data generated or analyzed during this study are included in this article. Further enquiries can be directed to the corresponding author.

## References

1. Antoni, L., et al., Intestinal barrier in inflammatory bowel disease. World J Gastroenterol, 2014. 20(5): p. 1165–79.

2. Vancamelbeke, M. and S. Vermeire, The intestinal barrier: a fundamental role in health and disease. Expert Rev Gastroenterol Hepatol, 2017. 11(9): p. 821–834.

3. Sheng, Y.H. and S.Z. Hasnain, Mucus and Mucins: The Underappreciated Host Defence System. Front Cell Infect Microbiol, 2022. 12: p. 856962.

4. Odenwald, M.A. and J.R. Turner, The intestinal epithelial barrier: a therapeutic target? Nat Rev Gastroenterol Hepatol, 2017. 14(1): p. 9–21.

5. Noah, T.K., B. Donahue, and N.F. Shroyer, Intestinal development and differentiation. Exp Cell Res, 2011. 317(19): p. 2702–10.

6. Neurath, M.F., D. Artis, and C. Becker, The intestinal barrier: a pivotal role in health, inflammation, and cancer. The Lancet Gastroenterology & Hepatology, 2025. 10(6): p. 573–592.

7. Soderholm, A.T. and V.A. Pedicord, Intestinal epithelial cells: at the interface of the microbiota and mucosal immunity. Immunology, 2019. 158(4): p. 267–280.

8. Soranno, D.E., et al., A review of gut failure as a cause and consequence of critical illness. Crit Care, 2025. 29(1): p. 91.

9. Rath, T., et al., Intestinal Barrier Healing Is Superior to Endoscopic and Histologic Remission for Predicting Major Adverse Outcomes in Inflammatory Bowel Disease: The Prospective ERIca Trial. Gastroenterology, 2023. 164(2): p. 241–255.

10. Orlemann, T., et al., Intestinal barrier healing is superior to transmural healing to prevent disease progression in clinical remittent patients with inflammatory bowel disease. Journal of Crohn‘s and Colitis, 2026. 20(1).

11. Bansil, R. and B.S. Turner, The biology of mucus: Composition, synthesis and organization. Advanced Drug Delivery Reviews, 2018. 124: p. 3–15.

12. Hansson, G.C., Role of mucus layers in gut infection and inflammation. Curr Opin Microbiol, 2012. 15(1): p. 57–62.

13. Overstreet, A.-M.C., et al., HMGB1 functions as a critical mediator of host defense at the gut mucosal barrier. Cell Host & Microbe, 2026. 34(2): p. 230–244.e13.

14. Tang, D., et al., The multifunctional protein HMGB1: 50 years of discovery. Nat Rev Immunol, 2023. 23(12): p. 824–841.

15. Bergstrom, K., et al., Proximal colon-derived O-glycosylated mucus encapsulates and modulates the microbiota. Science, 2020. 370(6515): p. 467–472.

16. Zhu, X., et al., Cytosolic HMGB1 controls the cellular autophagy/apoptosis checkpoint during inflammation. J Clin Invest, 2015. 125(3): p. 1098–110.

17. Mukherjee, P.K., et al., Stricturing Crohn‘s Disease Single-Cell RNA Sequencing Reveals Fibroblast Heterogeneity and Intercellular Interactions. Gastroenterology, 2023. 165(5): p. 1180–1196.

18. Daperno, M., et al., Development and validation of a new, simplified endoscopic activity score for Crohn‘s disease: the SES-CD. Gastrointest Endosc, 2004. 60(4): p. 505–12.

19. McCormick, D.A., L.W. Horton, and A.S. Mee, Mucin depletion in inflammatory bowel disease. J Clin Pathol, 1990. 43(2): p. 143–6.

20. Pullan, R.D., et al., Thickness of adherent mucus gel on colonic mucosa in humans and its relevance to colitis. Gut, 1994. 35(3): p. 353–359.

21. Kramer, C., et al., Ileal mucus viscoelastic properties differ in Crohn’s disease. Mucosal Immunology, 2024. 17(4): p. 713–722.

22. Barnich, N., et al., CEACAM6 acts as a receptor for adherent-invasive E. coli, supporting ileal mucosa colonization in Crohn disease. J Clin Invest, 2007. 117(6): p. 1566–74.

23. Gouyer, V., F. Gottrand, and J.L. Desseyn, The extraordinarily complex but highly structured organization of intestinal mucus-gel unveiled in multicolor images. PLoS One, 2011. 6(4): p. e18761.

24. Seifarth, C., M.E. Kreis, and J. Gröne, Indications and Specific Surgical Techniques in Crohn‘s Disease. Viszeralmedizin, 2015. 31(4): p. 273–9.

25. Sartor, R.B., Microbial Influences in Inflammatory Bowel Diseases. Gastroenterology, 2008. 134(2): p. 577–594.

26. Darfeuille-Michaud, A., et al., Presence of adherent Escherichia coli strains in ileal mucosa of patients with Crohn‘s disease. Gastroenterology, 1998. 115(6): p. 1405–13.

27. Small, C.-L.N., et al., Persistent infection with Crohn’s disease-associated adherent-invasive Escherichia coli leads to chronic inflammation and intestinal fibrosis. Nature Communications, 2013. 4(1): p. 1957.

28. Rhodes, J.M., R.R. Black, and A. Savage, Altered lectin binding by colonic epithelial glycoconjugates in ulcerative colitis and Crohn‘s disease. Dig Dis Sci, 1988. 33(11): p. 1359–63.

29. Robbe Masselot, C., et al., Human Fecal Mucin Glycosylation as a New Biomarker in Inflammatory Bowel Diseases. Inflamm Bowel Dis, 2023. 29(1): p. 167–171.

30. Kudelka, M.R., et al., Intestinal epithelial glycosylation in homeostasis and gut microbiota interactions in IBD. Nat Rev Gastroenterol Hepatol, 2020. 17(10): p. 597–617.

31. Jacobs, L.R. and P.W. Huber, Regional distribution and alterations of lectin binding to colorectal mucin in mucosal biopsies from controls and subjects with inflammatory bowel diseases. J Clin Invest, 1985. 75(1): p. 112–8.

